# Persistent post-COVID-19 smell loss is associated with inflammatory infiltration and altered olfactory epithelial gene expression

**DOI:** 10.1101/2022.04.17.488474

**Authors:** John B. Finlay, David H. Brann, Ralph Abi-Hachem, David W. Jang, Allison D. Oliva, Tiffany Ko, Rupali Gupta, Sebastian A. Wellford, E. Ashley Moseman, Sophie S. Jang, Carol H. Yan, Hiroaki Matusnami, Tatsuya Tsukahara, Sandeep Robert Datta, Bradley J. Goldstein

## Abstract

Most human subjects infected by SARS-CoV-2 report an acute alteration in their sense of smell, and more than 25% of COVID patients report lasting olfactory dysfunction. While animal studies and human autopsy tissues have suggested mechanisms underlying acute loss of smell, the pathophysiology that underlies persistent smell loss remains unclear. Here we combine objective measurements of smell loss in patients suffering from post-acute sequelae of SARS-CoV-2 infection (PASC) with single cell sequencing and histology of the olfactory epithelium (OE). This approach reveals that the OE of patients with persistent smell loss harbors a diffuse infiltrate of T cells expressing interferon-gamma; gene expression in sustentacular cells appears to reflect a response to inflammatory signaling, which is accompanied by a reduction in the number of olfactory sensory neurons relative to support cells. These data identify a persistent epithelial inflammatory process associated with PASC, and suggests mechanisms through which this T cell-mediated inflammation alters the sense of smell.

---

Recent studies have established that sustentacular cells of the OE in the nasal cavity are a main site of SARS-CoV-2 infection^1-3^. Transient olfactory loss, termed anosmia, occurs in a majority of subjects with COVID-19, but may persist following recovery^4-8^. The OE is the peripheral organ for olfaction, housing the primary olfactory sensory neurons, a barrier supporting cell layer termed sustentacular cells, and other populations including microvillar cells and basal stem or progenitor cells^9^ (**Figure 1a)**.

**Fig 1.**
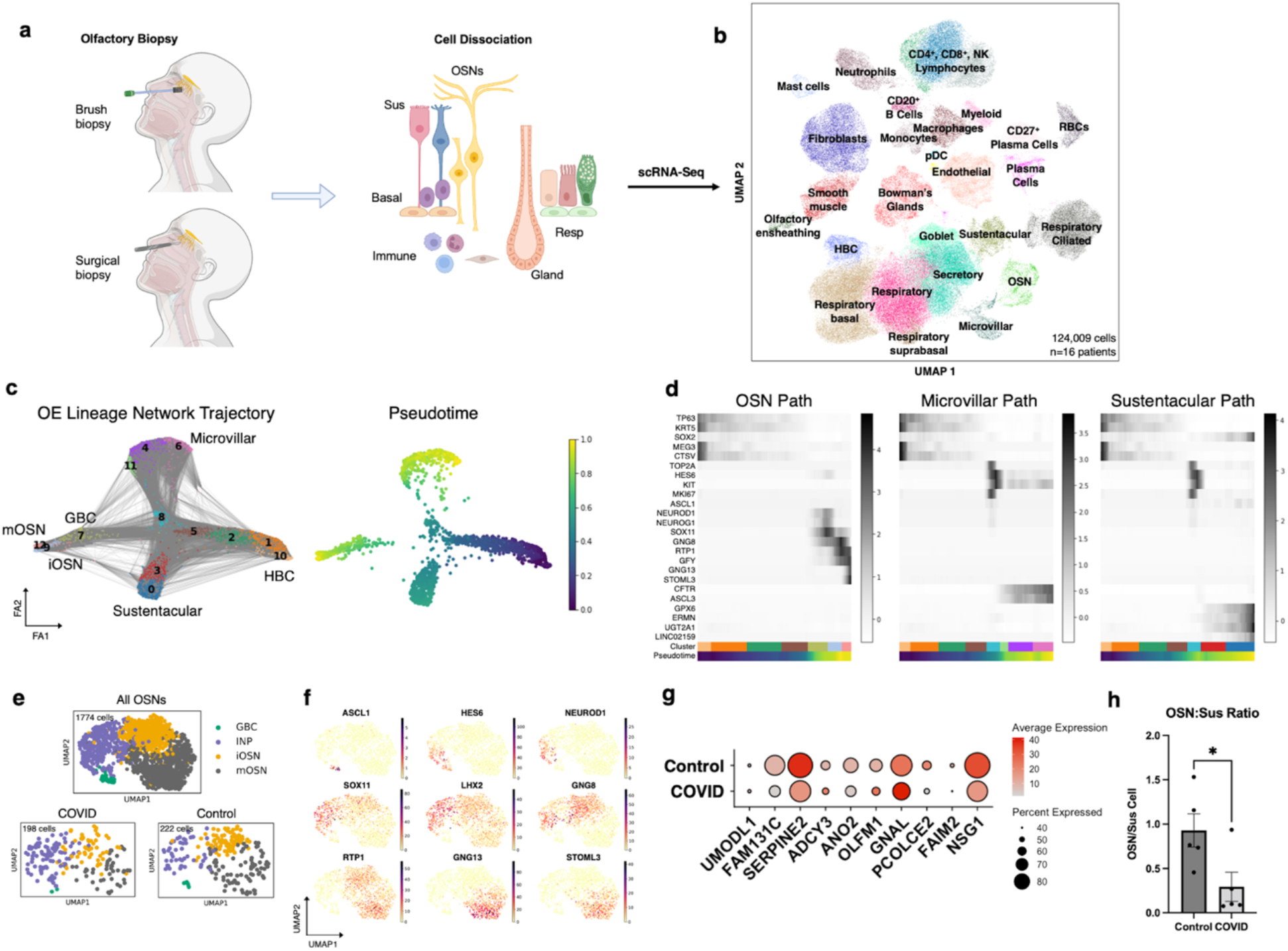
Nasal biopsies capture neurogenic OE populations and identify phenotypic alterations in PASC olfactory neurons. **a**, Schematic showing surgical or brush biopsy approach to obtain OE samples and generate dissociated cell suspensions for scRNA-seq. **b**, Visualization of combined PASC hyposmic and control biopsy datasets integrating n=16 human biopsies. **c**, Trajectory analysis; includes HBCs, OSNs, Sus cells, microvillar cells (n=7 biopsies, 4 PASC and 3 control). **d**, Heatmaps showing pseudotime progression for OE lineages; labeled based on UMAP relations in c; representative transcript markers for each stage along development are shown on Y-axis. **e**, UMAP showing annotations within the OSN cluster from b; “All OSNs” includes OSNs from 16 patients; “COVID” includes PASC hyposmic patients; “Control” includes 3 control normosmic biopsies. **f**, Gene expression plots markers along OSN differentiation pathway. **g**, Selected gene expression in PASC hyposmic (COVID) versus Control normosmic OSNs (mOSNs+iOSNs). **h**, Ratio of OSNs (mOSNs+iOSNs) to Sustentacular cells in each condition. Control n=5, COVID n=5; error bars indicate SEM; two-tailed t test, p=0.034.

The olfactory sensory neurons detect volatile odors via olfactory receptors (ORs) localized to the neuronal cilia at the nasal airspace^10^, and are replenished following damage by neurogenic basal cells^11-14^. Evidence from animal models suggests that acute transient anosmia may be due to sustentacular cell dysfunction or loss, and inflammation, both of which drive transient gene expression changes in olfactory neurons or alter the character of the mucus layer surrounding neuronal cilia^2,15^. In addition, polymorphisms in the UGT2A1/UGT2A2 locus, whose gene product is expressed in sustentacular cells, are associated with elevated risk of COVID-19-related loss of smell or taste^16^. With viral clearance, it is likely that normal epithelial reparative processes reconstitute the sustentacular cell population, restoring function^13^.

However, in the subset of subjects with lasting olfactory loss, what prevents recovery? There are several non-mutually-exclusive possibilities, including severe initial epithelial damage that eliminates the basal stem cell pools that normally are activated to reconstitute the neuroepithelium; infiltration of the OE by immune cell populations such that neuroinflammation or auto-immune phenomena perturb normal olfactory function and homeostasis through alterations in gene expression or other means; or central mechanisms that cause derangements in the olfactory bulbs of the brain or olfactory cortex. Evidence from animal models and human autopsies suggests that severe initial epithelial loss is unlikely, and that CNS involvement appears limited. However, no direct examination of olfactory tissue (including single cell RNA-sequencing) from humans suffering from PASC-olfactory dysfunction has been reported. Here, we obtained OE biopsies from subjects with lasting post-COVID olfactory loss, defined by objective olfactory testing, and examined samples using single cell RNA-sequencing (scRNA-seq) and immunohistochemistry to identify the cell populations and transcriptional alterations associated with PASC-olfactory dysfunction.

## RESULTS

We obtained nasal biopsy samples from subjects reporting olfactory dysfunction persisting at least 4 months since the onset of COVID-19 (**Figure 1a and Table S1**). Olfactory function was assessed with psychophysical testing (SIT, Smell Identification Test, Sensonics, Inc, Haddon Heights, NJ) confirming hyposmia. Most subjects also reported subjectively some component of parosmia, or distorted odor perceptions. Endoscopic olfactory mucosa biopsies were obtained either in the otolaryngology clinic, or in the operating room in subjects undergoing unrelated trans-sphenoidal procedures to access the pituitary for benign disease. Sinusitis or other known sinonasal disease was excluded by endoscopic exam and imaging, excluding bacterial infection, edema, polyposis. Biopsies were processed immediately for scRNA-seq analysis, as we have described previously^14,17^. Additional surgical olfactory samples were processed for histologic analysis, including a PASC hyposmic and a post-COVID normosmic (1.5-2 months post-COVID-19). Samples for scRNA-seq included 6 PASC hyposmics (age range 22-58 years; 5 female, 1 male, 4-16 months post-COVID-19 onset). For comparison, we analyzed 3 normosmic control samples (age range 51-71 years; 2 female, 1 male); to bolster these control data, we combined them with previously published datasets from normosmic and presbyosmic patients to generate an integrated single-cell sequencing dataset from a total of 16 subjects, permitting robust cluster annotation from >124,000 cells (**Figure 1b**). In a separate group of subjects, olfactory cleft mucus was collected from PASC hyposmics (n=15 subjects) or normosmic controls (n=13 subjects) for cytokine/chemokine assays.

### Nasal biopsies from control or PASC hyposmic subjects capture olfactory neuroepithelial cells

The OE is a self-renewing pseudostratified epithelium comprised of apical sustentacular and microvillar cells, mature OSNs, and immature OSNs emerging from basal stem and progenitor cells termed globose basal cells (GBCs) or horizontal basal cells (HBCs) (**Figure 1a**). Commonly, patches of respiratory epithelium are interspersed within the olfactory cleft region, comprised of secretory cells, ciliated cells and basal cells. Submucosal Bowman’s glands, vascular or stromal cells, and immune cell populations are abundant as well. To examine cell states and transcriptional profiles, we rapidly dissociated biopsies and processed live cell suspensions for scRNA-seq (**Figure 1b**). Uniform manifold approximation projection (UMAP) plots confirmed that the expected distribution of olfactory, respiratory and immune cells were captured for analysis. Importantly, we did not detect SARS-CoV-2 transcripts in scRNA-seq biopsy samples from any PASC hyposmics aligned to the viral reference genome; thus olfactory dysfunction in these patients is unlikely to reflect ongoing infection with SARS-CoV-2. Pseudotime analysis confirmed the expected OE lineage relationships and marker gene expression in the OE and in the neurogenic adult OE niche (**Figure 1c-f**).

### Olfactory neuron alterations are present in PASC hyposmic olfactory epithelium

Consistent with prior work exploring the acute effects of SARS-CoV-2 infection in human nasal epithelia, we observed changes in several transcripts associated with olfactory function in neuron clusters from PASC hyposmics compared to controls, including a reduction in the key adenylyl cyclase (ADCY3) that couples ORs to action potentials^15^ (**Figure 1g**). To quantify OSN numbers, we normalized OSN counts to sustentacular cell counts (**Figure 1h;** n=5 PASC hyposmic, 5 control, p=0.034, 2-tailed t test); we used this approach since OE biopsies can be variable and because there are often patches of respiratory-like metaplasia in biopsies. When normalized in this manner, OSNs were reduced relative to sustentacular cells in the PASC hyposmic samples compared to controls. However, despite the reduction in OSN number, in the PASC samples we observed no differences relative to control in the frequency of cells expressing ORs, the expression levels of the OR genes, or the distribution of ORs expressed across OSNs (**Figure S1**). Thus, the selective deficit in OSN numbers raises the possibility that PASC-related changes in immature and mature OSN cell number may underlie changes in smell.

### Olfactory sustentacular cells exhibit an immune response phenotype in PASC hyposmic subjects

Sustentacular cells express coronavirus entry genes, and SARS-CoV-2 has been found to infect this cell population during acute COVID-19^1,2,15^. Sustentacular cells have multiple functions as the apical barrier cell lining the OE, including detoxification of harmful chemicals via robust expression of biotransformation enzymes, modulation of the ionic content of the mucus layer in which OSN cilia are embedded, and feedback regulation of OE neuronal stem cells^18,19^. Based upon expression of canonical sustentacular markers such as ERMN^14^, CYP2A13, and GPX6, we identified 780 high quality sustentacular cells from PASC hyposmic or control samples (**Figure 2a**). We observed strong expression in sustentacular cells of UGT2A1, a gene shown via genome-wide association study (GWAS) to convey elevated risk of olfactory loss in COVID-19^16^. Differential gene expression (DE) analysis identified marked transcriptional alterations between PASC hyposmic or control sustentacular cells (**Figure 2b, S2**). Consistent with sustentacular cells mounting a response to inflammation, antigen presentation genes were markedly enriched in sustentacular cells derived from PASC samples (**Figure 2c and S2b,c)**, with minimal changes in typical markers of active viral infection such as CXCL10, PTX3, MX1 or CD46. Gene set enrichment analysis of the transcripts significantly upregulated in PASC hyposmics (log2 fold change >0.6, p<0.05) identified several biological processes including interferon signaling and antigen presentation (**Figure 2d**). Together, these findings suggest that sustentacular cells do not remain infected with SARS-CoV-2 during PASC, but rather appear to be responding to local pro-inflammatory cytokines in their micro-environment.

**Fig 2.**
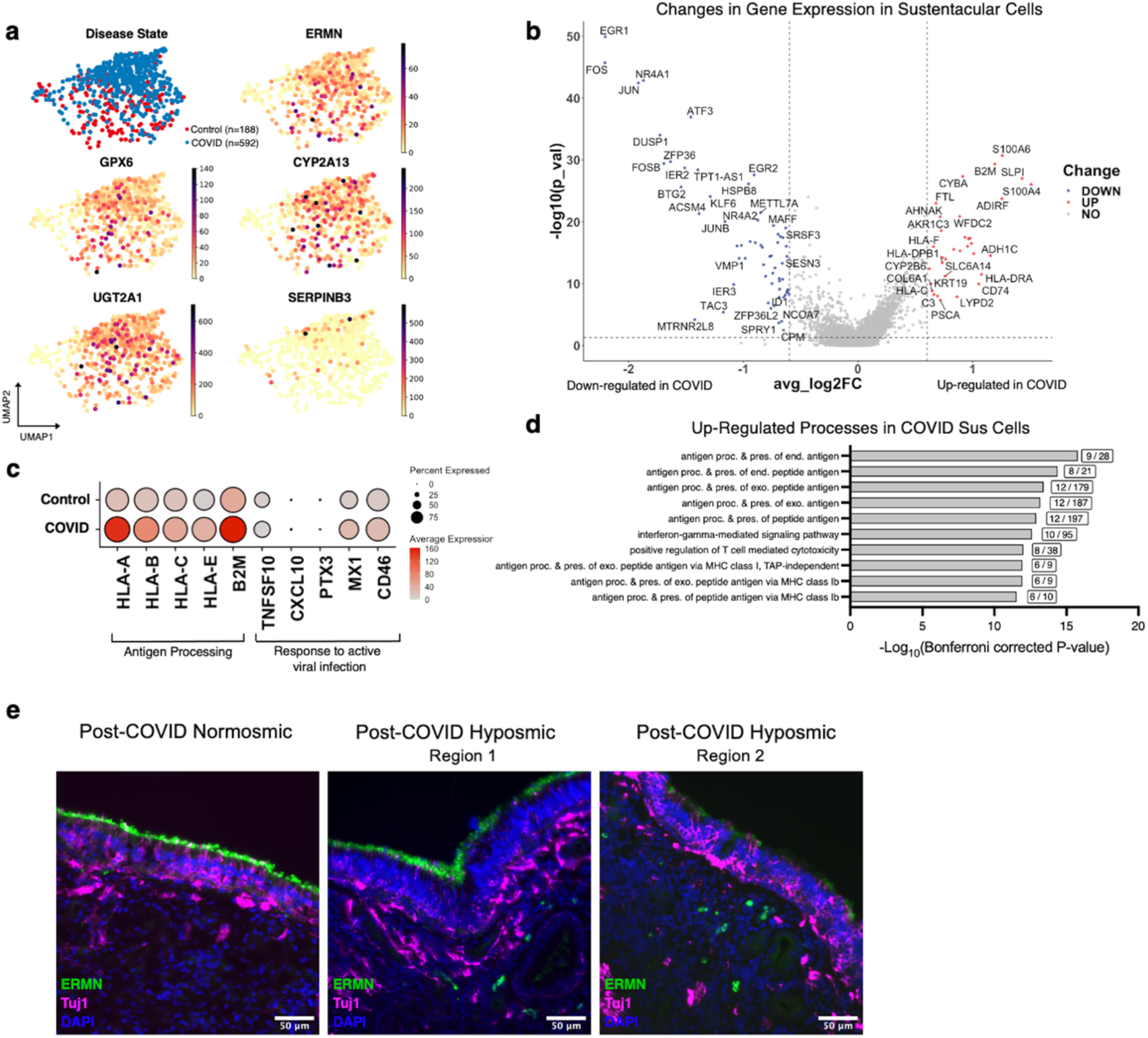
Sustentacular cell alterations persist in PASC hyposmic subjects. **a**, Gene expression visualization in PASC hyposmic and control sustentacular cell subset, n=9 biopsies. **b**, Differential sustentacular cell gene expression, PASC hyposmic versus controls; significant change indicates >0.6 log2 fold change, p<0.05. **c**, Visualization of selected antigen presentation genes and genes normally involved in responding to active viral infection in Control versus PASC hyposmic sustentacular cells. **d**, Gene set enrichment analysis for biological processes significantly upregulated in PASC hyposmic sustentacular (Sus) cells in b. **e**, Immunohistochemistry depicting representative labeling comparing sustentacular marker ERMN (green) and neuronal marker TUJ1 (magenta) in PASC hyposmic versus normosmic biopsies. Nuclei are stained with DAPI, bar=50 μm.

These findings were validated through immunohistochemical staining of a post-COVID normosmic biopsy and a post-COVID hyposmic biopsy (**Figure 2e**). Anti-ERMN selectively labels the apical region of sustentacular cells but not respiratory epithelial cells, showing strong uniform signal in the control (**Figure 2e**, left). TUJ1 antibody stains immature OSN somata and neurites, with abundant labeling in the control. Two representative regions in the hyposmic sample show abnormal labeling patterns, with one area showing intact sustentacular cells but few neurons (**Figure 2e**, middle) and another showing present but disorganized neurons and only weak patchy sustentacular label (**Figure 2e**, right).

### Specific T cell subpopulations are enriched in PASC hyposmic olfactory samples

The identification of sustentacular phenotypes harboring cytokine responsive transcriptional changes in PASC biopsies suggests the persistence of local immune cells providing inflammatory signaling. Accordingly, we explored all immune cell populations and identified the CD8^+^ and NK/NKT cell clusters as having robust expression of pro-inflammatory cytokines, specifically IFNG (**Figure 3a**). Using peripheral tissue lymphocyte markers^20^, we further subdivided these populations into NK/NKT, CD4, mixed CD4/CD8, CD8 effector (CD8 Teff), and CD8 tissue resident (CD8 Tres). We identified a specific CD8 Tres population, designated here as CD8 Tres *5*, with increased frequency in PASC hyposmics (**Figure 3b-d**, n=6 PASC, n=7 controls, p= 0.0015, 2-tailed t test). We next analyzed transcripts selectively enriched in this subpopulation, and their signaling ligands (**Figure 3e-g**). The genes enriched in this subset suggest that they are gamma-delta T cells, and a ligand-receptor model suggests potential interactions between signals — including interferon-gamma — from these T cells acting via specific receptors expressed by OSNs or sustentacular cells (**Figure 3e-g**).

**Fig 3.**
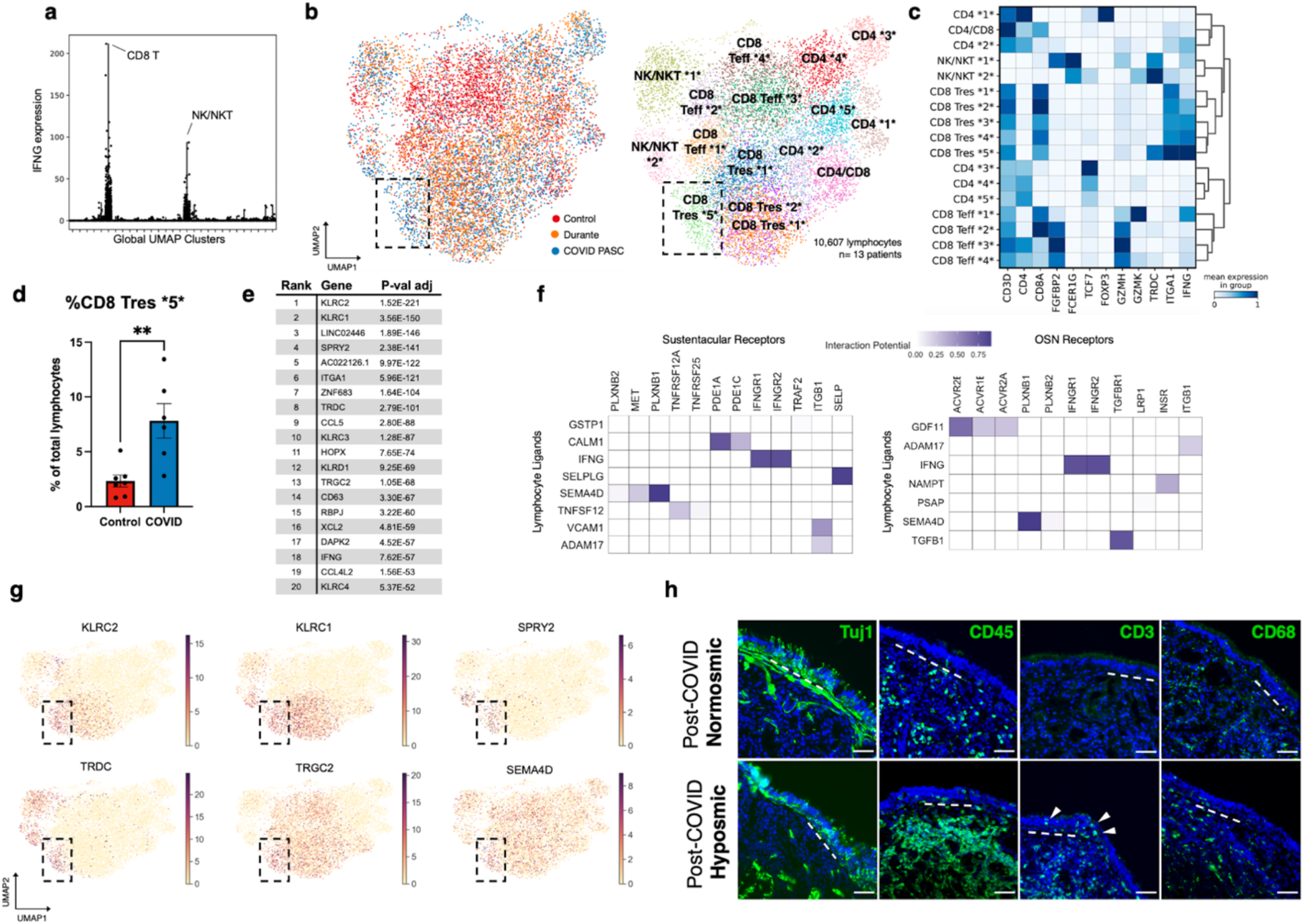
A lymphocyte subset is enriched in PASC hyposmic olfactory samples. **a**, Across all biopsy clusters, IFNG expression is highest in lymphocyte subsets, specifically in CD8^+^ and NK/NKT clusters. **b**, UMAP of all lymphocytes from CD8^+^, CD4^+^, and NK/NKT cell clusters, comparing PASC hyposmic (COVID) versus normosmic Controls; boxed area denotes cluster enriched in COVID PASC patients; cells from normosmic Durante et al (2020) dataset included for comparison. Annotations: Teff = effector T cells; Tres = resident T cells; subsets within categories are designated by numbers (i.e. CD4 *1* thru CD4 *5*). **c**, Selected gene expression and dendrogram clustering among lymphocyte subsets confirming annotations based on published marker genes; IFNG is enriched in Tres subsets, especially CD8 Tres *5*. **d**, Cell cluster designated CD8 Tres *5* is enriched in PASC hyposmics; control includes normosmic biopsies from “Control” and “Durante”; two-tailed T-test, p=0.0015. **e**, Top ranked (by adjusted p-value, Wilcoxon Rank-Sum with Bonferroni correction) transcripts enriched in CD8 Tres *5* cluster. **f**, NicheNet analysis of CD8 Tres-derived ligands and their receptors in either sustentacular cells or OSNs, depicting interaction potential. **g**, Selected gene expression plots of statistically significantly enriched genes in PASC hyposmic-specific cluster CD8 Tres *5* (boxed area), including gamma-delta T cell markers and SEMA4D, identified by NicheNet analysis. **h**, Representative immunohistochemistry from normosmic post-COVID biopsy versus PASC hyposmic biopsy; TUJ1 neuronal label appears more prominent in normosmic; hyposmic has dense CD45^+^ immune infiltration, with prominent CD3^+^ lymphocytic infiltration, which is absent in the normosmic sample; arrowheads indicate lymphocytes within the OE; dashed lines mark basal lamina; scale bars = 50 μm.

In a separate cohort of PASC hyposmics and controls, olfactory mucus was assayed to measure cytokines and chemokines (**Figure S3**). Consistent with a lack of severe cytotoxic inflammation, we verified that there were no markedly elevated changes in IL-1≻or TNFα? We confirmed the presence of IFN-γ, although no significant difference was apparent in mucus, despite differences in IFN-γ secreting cells identified by scRNA-seq. Finally, we qualitatively confirmed the T cell infiltrates observed by sc-Seq through immunohistochemistry on biopsies from control or post-COVID-19 hyposmic subjects (**Figure 3h**). The COVID hyposmic staining exhibits dense CD45^+^ immune infiltration, and prominent CD3^+^ lymphocytic infiltration, both of which are absent in the normosmic. CD68 myeloid labeling pattern appears qualitatively similar in both conditions. Together, we identify in PASC hyposmic OE evidence of a persistent lymphocytic response that may provide signaling sufficient to drive localized IFN-response phenotypes in sustentacular cells. This observation suggests a coupling between immune infiltration and sustentacular cell gene expression that could influence olfactory function.

## DISCUSSION

Understanding PASC pathophysiology is a global priority, as lasting dysfunction following SASR-CoV-2 infection may involve multiple systems, including olfactory, neuropsychiatric, respiratory, or cardiovascular^21,22^. Current evidence for COVID-19 damage within the human OE comes largely from autopsy studies from subjects who died from severe *acute* COVID-19^3,15^. These studies lack objective measurements of smell, and samples were obtained in the context of significant prior medical intervention. While those studies provide valuable insights into the cell types infected by SARS-CoV-2 with the OE, the mechanisms involved in lasting PASC olfactory dysfunction have remained elusive. We provide here the first analysis of olfactory biopsies from PASC hyposmic humans. Our efforts here focused on the subset of subjects with olfactory dysfunction lasting at least 4 months following COVID-19 as documented by objective olfactory testing. Our results comparing scRNA-seq data between endoscopically-guided OE biopsies from PASC hyposmics and control normosmic samples suggest a model in which altered immune cell – OE interactions drive functional changes in sustentacular cells and olfactory sensory neurons.

Our findings are consistent with a model in which ongoing indolent localized immune cell responses drive phenotypic changes in sustentacular cells and OSNs; the changes observed in OSNs (including a reduction in the expression of key signaling intermediates and a relative reduction in cell number) could explain sensory dysfunction including hyposmia or parosmia. Major findings identified here include OSN transcriptomic changes that align with some of the acute findings identified in autopsy subjects^15^, suggesting a persistence of an underlying non-cell autonomous signal. A likely identity for this signal emerges from the demonstration here of interferon response signatures in the sustentacular cells, along with the presence of local lymphocyte populations unique to the PASC OE samples expressing IFN and CD8 Tres markers. Both sustentacular cells and HBCs modulate OE renewal and homeostasis via multiple signaling pathways^13^. Importantly, in mouse models^23^ and in presbyosmic humans^17^ HBCs exhibit immune responsive phenotypes, including signaling interactions that can recruit additional immune cells.

It is also interesting to compare the phenotypes observed here in humans with PASC-related smell loss and those observed previously in hamsters acutely infected with SARS-CoV-2^2,15^. In the hamster model, a wide array of immune cells (including macrophages, neutrophils and monocytes) infiltrates the epithelium in the first several days after infection, before resolving nearly completely within two weeks. Our observation of a persistent infiltration of T cells in the human OE months after infection suggests that PASC patients may have a selective immunological response to prior infection that differs from the immunological responses generated acutely.

Thus, our data are consistent with a provisional model in which a dysregulated immune cell – HBC/sustentacular cell – OSN axis arises in the PASC OE, with a resultant sensory dysfunction. How and why this occurs in a subset of patients remains to be defined, but analysis of macrophages in COVID-19 subjects has shown that acute SARS-CoV-2 infection drives a pro-inflammatory reprogramming that is thought to induce long-term alterations in other immune cell function^24^. Furthermore, unresolved immune responses, including activation of specific CD8^+^ T cell clonotypes during convalescence following SARS-CoV-2 infection, have been reported^25^. It is tempting to speculate that a similar process may initiate the local OE immune cell alterations identified in our PASC hyposmic samples.

These data also are relevant to several alternative mechanistic hypotheses about how SARS-CoV-2 might cause causing long term olfactory loss. One possibility is that severe initial widespread cell damage might overwhelm the capacity of basal stem cells to reconstitute the epithelium, but our samples suggest that at least many areas of the olfactory cleft harbor intact OE comprised by OSNs, sustentacular cells and basal cells. Persistent viral infection could also drive ongoing damage, but we find no evidence for active SARS-CoV-2 infection in our samples. Another possibility is that anosmia/parosmia is the consequence of severe ongoing mucosal inflammation, but our patients did not exhibit clinical inflammatory findings of local edema, polyposis or infection, and the molecular signatures identified in the OE were not consistent with broad inflammatory responses. It is important to note that central mechanisms may contribute to PASC-related smell loss, warranting further study; however, there is little evidence for SARS-CoV-2 infection of neurons in humans^3^,and at least some of the observed imaging changes in the olfactory bulb or cortex^26^ could reflect reduced peripheral input due to OE damage (the clear site of viral infection), or diffusion of inflammatory intermediates across the cribriform plate.

There are several caveats related to our conclusions. Given challenges related to the pandemic, it has been difficult to obtain samples from large numbers of patients, and thus our conclusions are driven by findings we observe in common across our limited set of patients; furthermore, in these studies we have merged patient samples obtained by two different methods (surgical excision and brush biopsy). Finally, although we were careful to obtain biopsies from within the olfactory cleft region in these experiments, the possibility of sample-to-sample variation in the specific contents of each OE biopsy in unavoidable.

The pandemic has highlighted the unmet need for new effective treatments for olfactory loss. The mechanistic insights provided here suggest potential new therapeutic strategies. For instance, selectively blocking local pro-inflammatory immune cells or directly inhibiting specific signaling nodes may interfere with a loop disrupting OE homeostasis or repair. The location of the OE, lining the olfactory cleft in the nose, is amenable to localized topical drug delivery, which may provide a means to avoid systemic or off-target effects of novel therapeutic agents. Further studies testing therapeutics in animal models and humans, and longer follow up with PASC olfactory dysfunction patients, will provide ongoing insights regarding the etiology and management of olfactory sensory dysfunction.

## METHODS

### Biopsy Collection

All biopsy samples reported here were collected under Duke University School of Medicine IRB protocols 00088414 and 00105837. Patients were administered the Smell Identification Test (SIT) prior to tissue collection to assess olfactory function. For surgical biopsies, olfactory mucosa was collected either in the operating room in patients undergoing transsphenoidal surgery for resection of a benign pituitary tumor or in the clinic (3 subjects). Briefly, using endoscopic visualization, olfactory cleft mucosa was sharply incised and elevated from underlying bone, and then excised with a through-cutting ethmoid forceps. For nasal cytology brush biopsies (3 subjects), tissue was collected in clinic by gently positioning a cytology brush (Cat#4290, Hobbs Medical Inc, Stafford Springs, CT) in the olfactory cleft under endoscopic visualization (**Figure S4**). The brush was rotated briefly to collect surface mucosal cells. In all cases, samples were placed into collection solution (Hanks’ Balanced Salt Solution (HBSS) or Hibernate E medium, with 10% fetal bovine serum (FBS; all from Thermo Fisher, Waltham, MA) on ice and processed immediately for analysis.

### Biopsy Processing

Surgical biopsy tissues were divided into smaller pieces sharply. All biopsies were digested for 15 minutes at 37°C with an enzyme cocktail comprised of Dispase/Collagenase A/EDTA mix, 2mg/mL Papain, and DNAse I (all from StemCell Tech, Vancouver, BC, Canada) with frequent gentle trituration. After 15 minutes, Accutase (StemCell Tech) was added, and samples were incubated for an additional 5 minutes at 37°C. At the end of 5 minutes, FBS was added. If samples still contained large pieces of tissue, they were filtered through a 250µm filter. All samples were then filtered through a 70µm filter and centrifuged 5 minutes at 400 x g. If abundant red blood cells were observed in the pellet, tissues were re-suspended in ACK Lysing Buffer (Thermo) and incubated at room temperature for 3-5 minutes while gently rocking. Samples were washed, spun and resuspended in HBSS or Hibernate-E containing non-acetylated bovine serum albumin 1 mg/ml (Thermo), anti-clumping reagent 0.5 µl/ml (Gibco), and N-acetyl cysteine 5 µg/ml (Sigma-Aldrich, St. Louis, MO) to a final concentration of 1 million cells/mL.

### Single-Cell Sequencing

Samples were processed for single cell analysis as described previously^17^. Briefly, cells were quantified with a viability stain on an automated counter (Cellaca MX, Nexcelom) and loaded onto a Chromium controller (10X Genomics, Pleasanton, CA) for cell capture and bar coding targeting 10,000 cells, per the 3’ v3.1 gene expression protocol per manufacturer’s instructions. Reverse transcription, amplification, library preparation, and sequencing (NovaSeq, Illumina) were performed per protocol.

### Single-Cell RNA-Seq Analysis

Illumina base call files were converted to FASTQ files and processed through CellRanger Counts 6.1.2 (10X Genomics), aligned to either a human reference genome (GRCh38) or a combined reference genome containing human and SARS-CoV-2 genomes^27^.

Starting from the raw cell by gene count matrices, data integration and preprocessing were performed using Scanpy (v1.8.2) and scvi-tools (v0.15.2). For accurate cell type identification, the data generated in this study were combined with our published human olfactory datasets^14,17^. Highly-variable genes (HVGs) were identified using the scvi-tools “poisson_gene_selection” function (with patient id as the batch key), and the raw counts for these gene subsets were used as the input to the variational autoencoder (scVI) model. An scVI model (using the top 3000 HVGs) was trained for 500 unsupervised epochs with the default learning rate (with early stopping when the ELBO validation metric did not improve for at least 20 epochs) with the default parameters (10 latent dimensions, 128 nodes per hidden layer), a negative binomial observation model (gene_likelihood=“nb”), the percentage of mitochondrial genes as a continuous covariate, and categorical covariate keys for the patient condition and patient id categorical variables (which thus performed dataset integration and batch correction for the purposes of cell type identification). A k-nearest neighbor graph was constructed from the resulting 10-dimensional latent embedding (using k=15 neighbors). The knn graph which was used for cell type clustering via the Leiden algorithm (resolution=1.2) and as the input to the UMAP algorithm (with min_dist=0.5) for visualization. Clusters of dying cells containing high percentages of mitochondrial genes and low total counts as well as a cluster of cell doublets were removed, and the above procedure starting from the HVG identification was repeated (but with Leiden clustering resolution=1.6). The resulting cell type clusters were merged and manually annotated based on known cell type markers.

After identifying and annotating the broad clusters, cell types of interest were further subclustered in an iterative manner, using the same scVI embedding approach, starting from the reidentification of HVGs for each subset. After training a scVI model using only cell types in the olfactory epithelium (including olfactory HBCs, sustentacular cells, Bowman’s gland cells, microvillar cells, and OSNs), an additional cluster of OSN-sustentacular cell doublets was identified and removed. Next, an scVI model was trained on the OSNs and microvillar cells (except using 2000 HVGs and 100 hidden nodes in the scVI model), and the resulting clusters from this model identified cell types of the OSN lineage; a final scVI model (using 2000 HVGs and 100 hidden nodes) was used to embed and cluster these cells and the OSN lineage cell types were manually annotated using known markers for GBCs (TOP2A, ASCL1), INPs (NEUROD1, SOX11), iOSNs (GAP43, DCX, GNG8), and mOSNs (GNG13, STOML3). The same approach was also used to further subcluster the broad lymphocyte cluster that contained the CD4^+^ T cells, CD8^+^ T cells and NK cells starting from the top 2000 HVGs from these cells and the resulting scVI embedding was then clustered (resolution=1.1) to identify the lymphocyte subtypes.

Trajectory and pseudotime analyses were performed using OSN lineage cell types identified in the second iteration of the scVI model trained on these cells. Bowman’s gland cells were excluded from analysis. A new neighborhood graph was computed using n_neighbors=100 and n_pcs=20, and cells were reclustered using the default leiden algorithm with resolution=1.5. Cluster connectivity was then calculated using partition-based graph abstraction (PAGA) with default settings. PAGA plots were constructed using threshold=0.2. For plotting of pseudotime heatmaps, leiden clusters were ordered based on PAGA connectivity predictions.

Transcriptome distances were calculated from the pairwise correlation distance matrix of the embedding in the 10-dimensional scVI latent space embedding for cells from the OSN lineage. Transcriptome distances were summarized for each OSN cluster -condition (control vs PASC) pair by taking the median pairwise transcriptome distance between cells of each pair.

For additional plots, such as differential expression analysis and further subsetting and analysis of immune cell populations, filtered outputs were analyzed in R (v4.1.1) using the Seurat toolkit (v4.1.0)^28^. Processed anndata objects from Scanpy were converted to R objects preserving all meta data (including scVI clusters) using the LoadH5Seurat function from SeuratDisk. Data were normalized using relative counts normalization prior to differential expression analysis. Differentially expressed genes were found using the FindMarkers function with default settings (Wilcoxon Rank-Sum) and plotted using ggplot2 (identifying significant DE genes with >log2 fold-change, adjusted p<0.05). Cluster markers of lymphocyte subsets were identified using FindAllMarkers with default settings.

NicheNet analysis was conducted in R with the nichenetr package (V 1.1.0) using the default ligand-target prior model, ligand receptor network, and weight integrated networks^29^. Specifically, cell populations of interest (i.e. lymphocyte clusters, OSNs, and sustentacular cells) with normalized gene expression were subset out from processed R objects (from anndata) and used as input for the appropriate receiver and sender populations.

### Immunohistochemistry

Samples for histology were collected in Hanks’ Balanced Salt Solution (HBSS, Gibco) + 10% FBS. Tissues were fixed with 4% paraformaldehyde (Sigma, St. Louis) in phosphate buffered saline (PBS) for 4 hours at room temperature. Samples were washed with PBS and then incubated on a rocker at 4°C for 5-7 days in 30% sucrose, 250mM EDTA, and PBS. Samples were then flash frozen in OCT compound (VWR, Radnor, PA), sectioned at 10µm on a cryostat (CryoStar NX50, Thermo Fisher) and collected on Superfrost plus slides (Thermo Fisher).

Tissue sections were rehydrated in PBS and blocked in 5% normal goat serum in PBS with 0.1% Triton X-100. Anti-Tubulin β3 (BioLegend, clone TUJ1, 1:500), Anti-CD45 (BioLegend, clone HI30, 1:100), Anti-CD3 (BioLegend, clone HIT3a, 1:100), Anti-CD68 (BioLegend, clone BL13756, 1:100) or anti-ERMN (Thermo, #PA5-58327, 1:500) primary antibodies diluted in blocking buffer were incubated on tissue sections for 1 hour at room temperature or overnight at 4 degrees. Following PBS washes, tissues were incubated with fluorescent conjugated secondary antibodies for 45 minutes (Jackson ImmunoResearch, West Grove, PA). Vectashield with DAPI (Vector Laboratories, Burlingame, CA) was applied to each section prior to coverslip. All images were acquired on a Leica DMi8 microscope system (Leica Microsystems). Images were analyzed using ImageJ software (V 2.3.0), and scale bars were applied using metadata from the original Leica acquisition software files.

### Olfactory mucus assays

Mucus was obtained from the olfactory cleft using absorbant filter paper under endoscopic guidance per an approved IRB protocol at UC San Diego (#210078). Cohorts included PASC hyposmics (n=15 patients) or control normosmics (n=13), based on psychophysical testing using the SIT. A fluorescent bead-based multiplex assay (LegendPlex, Biolegend) was used to quantify 13 cytokines/chemokines through flow cytometry.

### Statistics

All sequencing data set analyses were performed in Python or R using the toolkits and packages described above. Plots were produced using Scanpy, or ggplot2 in associated R toolkits^30^, or Graphpad Prism 9. Cell phenotype comparisons between PASC hyposmic and control samples were performed using unpaired 2-tailed t-test, with significance defined as p<0.05. Error bars represent standard error of the mean. DE gene sets were analyzed for gene ontology, cellular pathway or tissue output terms using ToppGene Suite^31^.

#### Study Approval

All human subjects studies were performed under protocols approved by Institutional Review Boards.

#### Disclosures

BJG discloses advising for Mirodia Therapeutics and consulting for Frequency Therapeutics.

## Acknowledgements

We thank the patients who generously agreed to provide biopsy samples for this research. We appreciate the expert technical assistance of the Duke Molecular Genomics Core, and bioinformatics assistance from Vaibhav Jain. We also thank Clinical Research Coordinators, Amy Walker and Victoria Eifert, for expert assistance. Graphical schematics were created with Biorender.com.

## Author Contributions

Conceptualization: B.J.G; Methodology: J.B.F., R.G., A.O., D.J., R.A-H., S.S.J., C.H.Y.; Data analysis: B.J.G., J.B.F., T.K., S.A.W., E.A.M., S.R.D., D.H.B., T.T.; Writing or manuscript review: J.B.F., B.J.G., H.M., D.H.B., T.T., S.R.D.; Funding acquisition: B.J.G.

## Data Availability

Data sets from scRNA-seq are deposited in GEO (accession numbers will be available upon publication).

## Code Availability

Any requests regarding code used here should be directed to the corresponding author.

## SUPPLEMENTAL FIGURES

**Table S1.** Biopsy samples processed for scRNA-seq. Immunohistochemistry (IHC); single cell RNA-sequencing (scRNA-seq).

|  | Sample | Age | Sex | UPSIT | Months Post-COVID | Type of Biopsy | Assay |
| --- | --- | --- | --- | --- | --- | --- | --- |
| **Control Normosmic** | Control 1 | 71 | F | 37 | - | Surgical | scRNA-Seq |
|  | Control 2 | 51 | M | 37 | - | Surgical | scRNA-Seq |
|  | Control 3 | 66 | F | 34 | - | Surgical | scRNA-Seq |
|  | Control 4 | 43 | M | - | 2 | Surgical | IHC |
| **COVID Hyposmic** | COVID 1 | 28 | F | 17 | 6 | Surgical | scRNA-Seq |
|  | COVID 2 | 31 | M | 28 | 4 | Surgical | scRNA-Seq |
|  | COVID 3 | 25 | F | 17 | 6 | Surgical | scRNA-Seq |
|  | COVID 4 | 58 | F | 24 | 13 | Brush | scRNA-Seq |
|  | COVID 5 | 22 | F | 29 | 13 | Brush | scRNA-Seq |
|  | COVID 6 | 58 | F | 24 | 16 | Brush | scRNA-Seq |
|  | COVID 7 | 29 | F | 23 | 1.5 | Surgical | IHC |

**Fig. S1.**
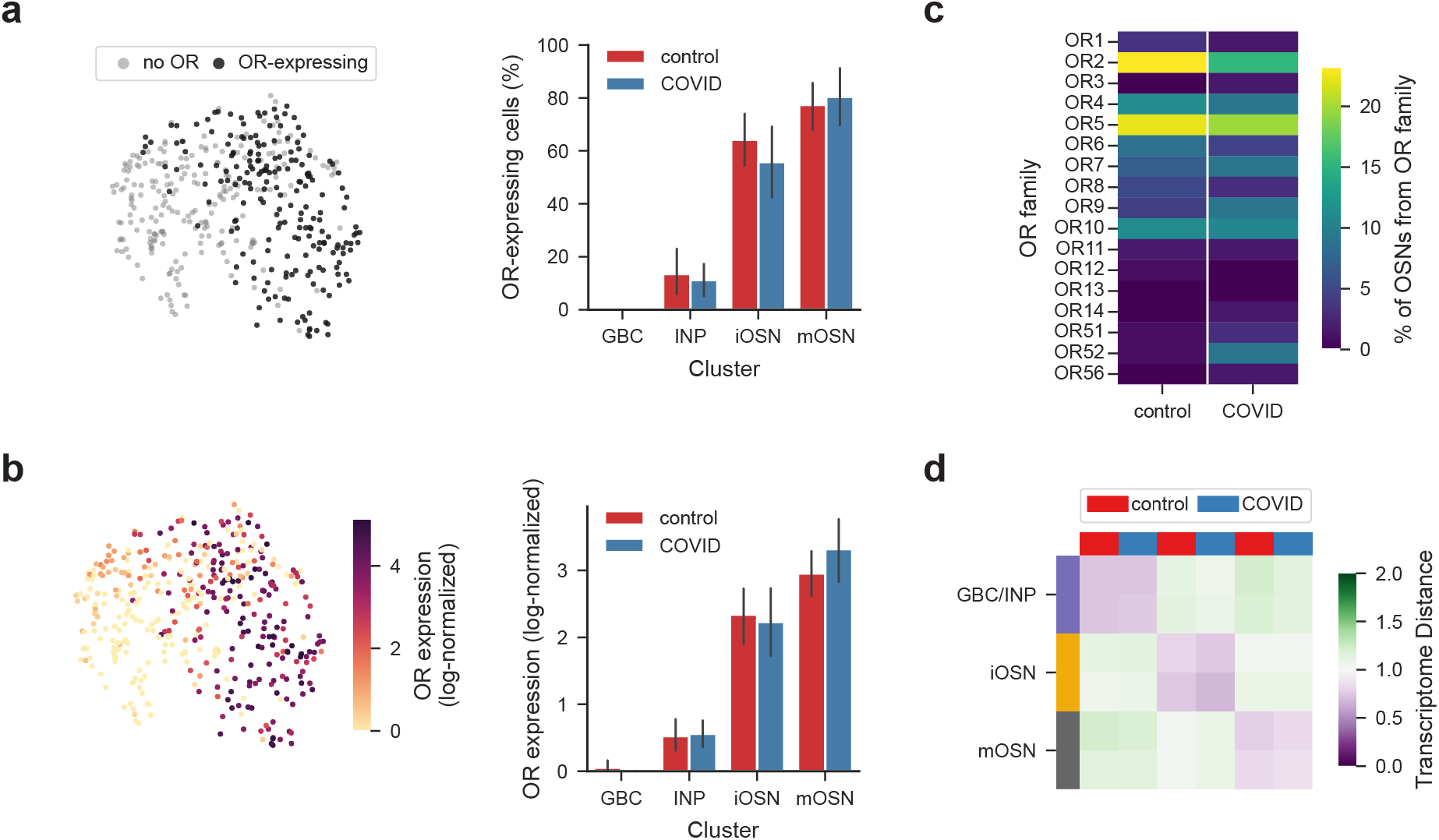
Olfactory receptor expression dynamics are unaltered in PASC olfactory neurons. **a**, (left) UMAP representation of the developing OSN lineage, colored by OR expression. (right) The percentage of cells expressing ORs for each cluster, for cells from normosmic control and hyposmic PASC patients. Error bars depict the mean depict the mean and the bootstrapped 95% confidence intervals of the mean. **b**, As in (a), except for the log-normalized expression level of the top OR expressed in each cell. **c**, Percent of OSNs whose expressed OR was from each OR family, for OR-expressing cells from normosmic controls and hyposmic PASC biopsies. **d**, Transcriptome distances (see methods) between pairs of cells from either normosmic or PASC patients from each OSN cluster. Pairs of cells from the same cluster from either condition are more similar to each other than those from the same condition from different clusters, suggesting that developing and mature neurons from PASC patients are similar to those of normosmic controls.

**Fig. S2.**
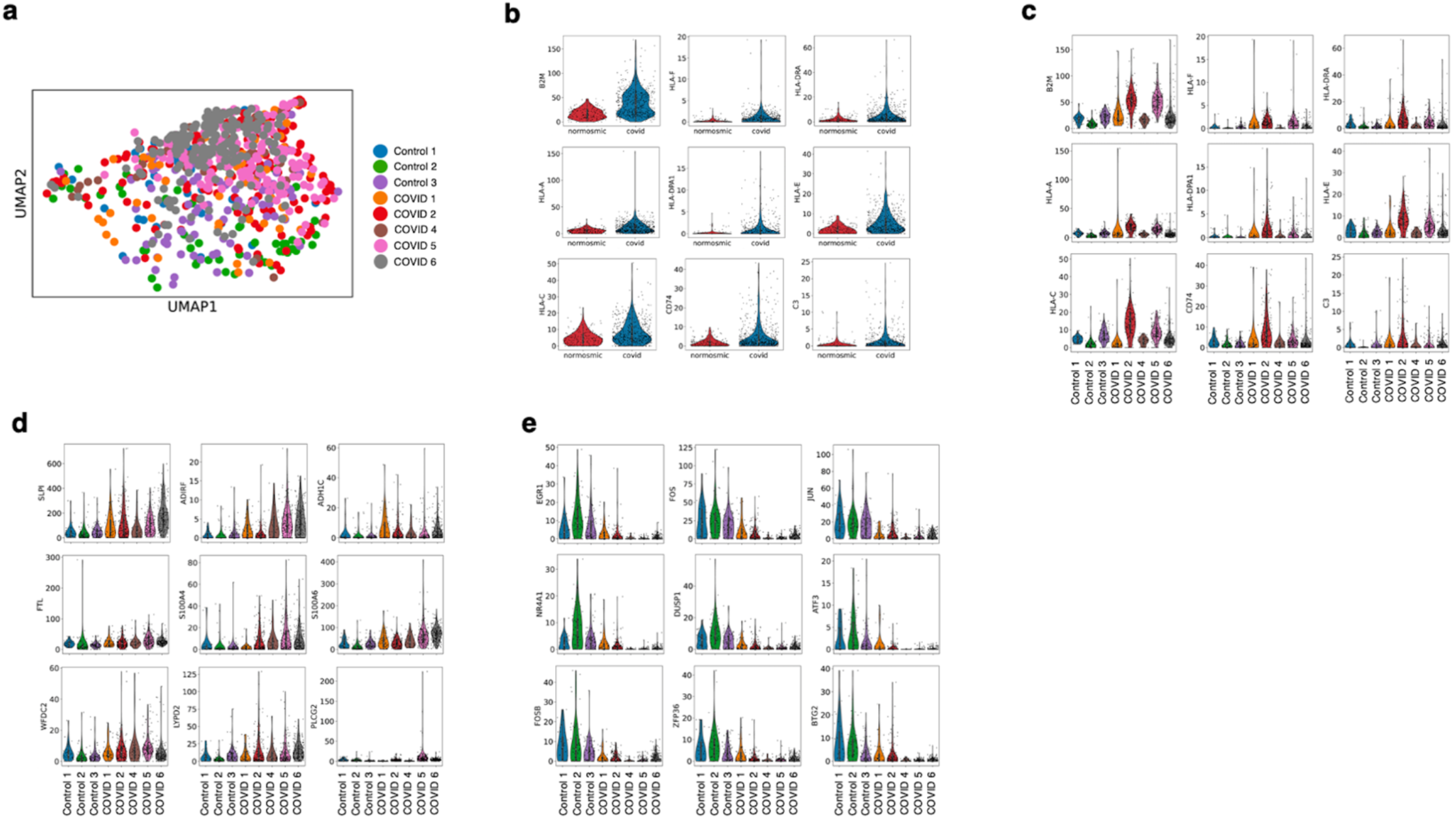
Interferon response pathways across PASC hyposmic sustentacular cells. **a**, UMAP of sustentacular cluster shows integration of patients without batch effects. Note in legends that biopsy from subject “COVID 3” lacks sustentacular cells and is therefore omitted. Statistically significantly up-regulated genes related to antigen presentation identified on volcano plot in Fig. 2b, separated by condition (**b)** and origin patient (**c)** show that these genes are up-regulated across multiple COVID patients. **d and e**, Other statistically significant up-regulated (d) and down-regulated (e) genes in COVID sustentacular cells unrelated to antigen presentation.

**Fig. S3.**
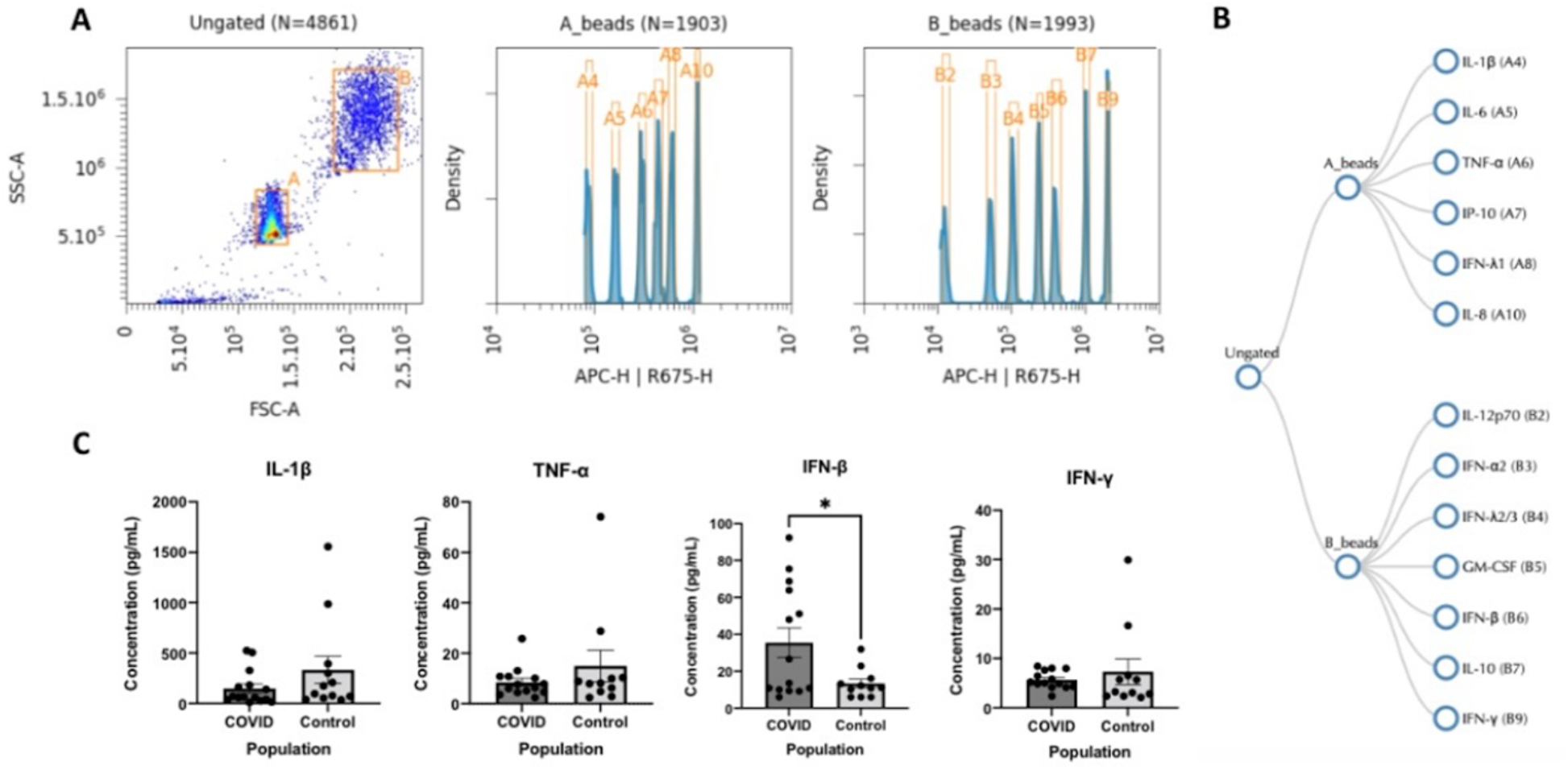
Olfactory mucus assays. Mucus was obtained from the olfactory cleft using absorbant filter paper under endoscopic guidance from PASC hyposmics (n=15 patients) or control normosmics (n=13). Quantification of cytokines/chemokines reveals only subtle changes rather than marked inflammation. Note that IFNs are detected in mucus at low levels, consistent with localized expression as identified via separate RNA-seq approaches.

**Fig. S4.**
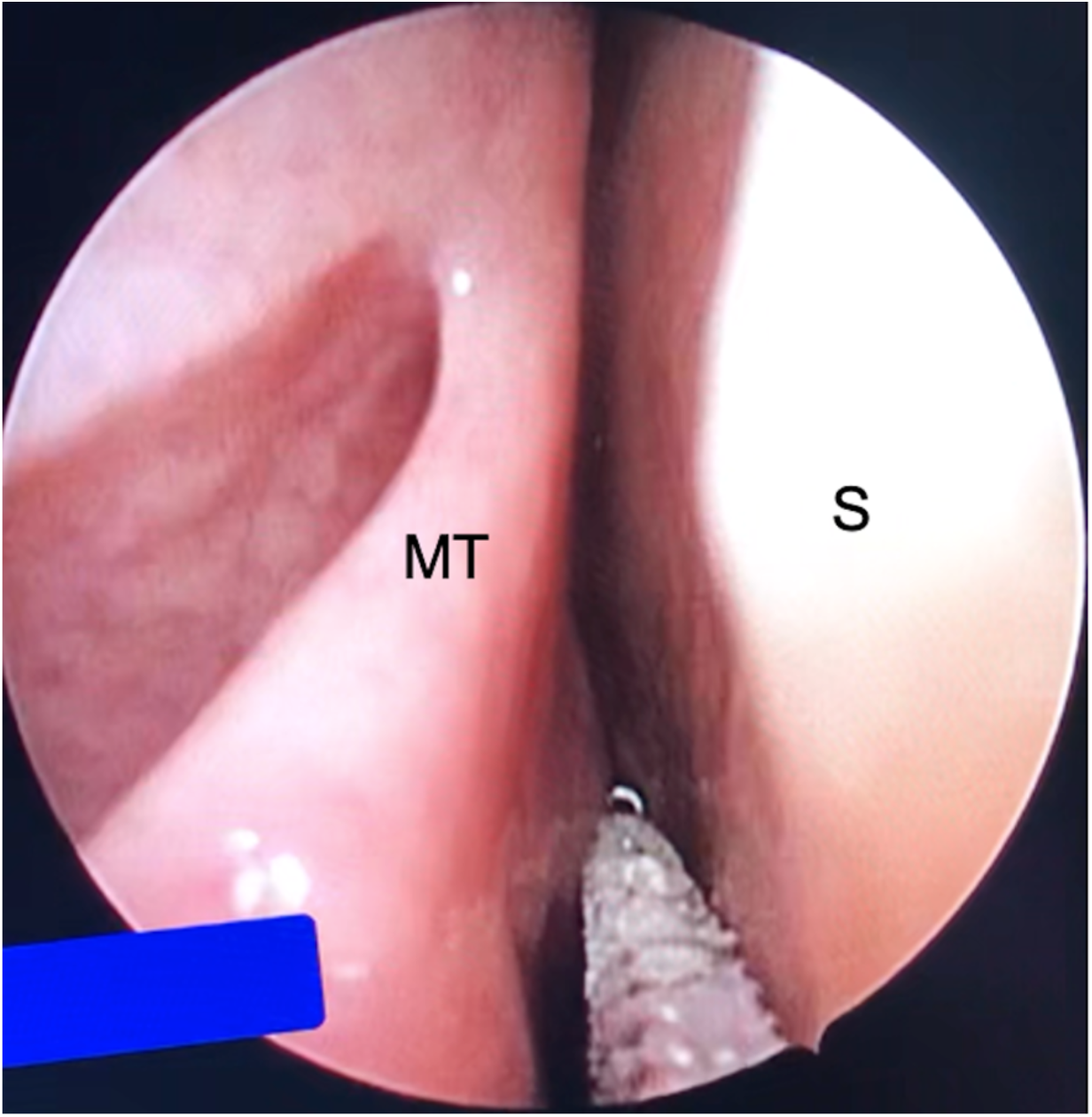
Endoscopic image showing biopsy of olfactory cleft region using cytology brush technique. Here, the brush is in place along the right nasal septum (S) superiorly, medial to the middle turbinate vertical lamella (MT) and positioned posteriorly approaching the superior turbinate.

